# Molecular mechanism of depolarization-dependent inactivation in W366F mutant of Kv1.2

**DOI:** 10.1101/316174

**Authors:** H. X. Kondo, N. Yoshida, M. Shirota, K. Kinoshita

## Abstract

Voltage-gated potassium channels play crucial roles in regulating membrane potential. They are activated by membrane depolarization, allowing the selective permeation of potassium ions across the plasma membrane, and enter a nonconducting state after lasting depolarization of membrane potential, a process known as inactivation. Inactivation in voltage-activated potassium channels occurs through two distinct mechanisms, N-type inactivation and C-type inactivation. C-type inactivation is caused by conformational changes in the extracellular mouth of the channel, while N-type inactivation is elicited by changes in the cytoplasmic mouth of the protein. The W434F-mutated Shaker channel is known as a nonconducting mutant and is in a C-type inactivation state at a depolarizing membrane potential. To clarify the structural properties of C-type inactivated protein, we performed molecular dynamics simulations of the wild-type and W366F (corresponding to W434F in Shaker) mutant of the Kv1.2-2.1 chimera channel. The W366F mutant was in a nearly nonconducting state with a depolarizing voltage and recovered from inactivation with a reverse voltage. Our simulations and 3D-RISM analysis suggested that structural changes in the selective filter upon membrane depolarization trap potassium ions around the entrance of the selectivity filter and prevent ion permeation. This pore restriction is involved in the molecular mechanism of C-type inactivation.

## INTRODUCTION

Voltage-gated potassium (Kv) channels conduct K^+^ ions selectively in response to membrane depolarization and regulate the amplitude and duration of action potentials. They are homotetramers of subunits containing six transmembrane helices named as TMH1 through TMH6; TMH1–TMH4 form the voltage sensor domain (VSD) and TMH5–TMH6 form the pore domain, which is composed of a central cavity (CC, inner mouth) and selectivity filter (SF). Ion permeation through the pore domain is dynamically regulated by several types of gating mechanisms (1). Upon depolarization of membrane potential, positively charged TMH4s of the VSD move toward the extracellular side, leading to channel opening (activation), whereas they move towards the intracellular side when the membrane potential returns to the resting level, leading to channel closure (deactivation). Although this activation-deactivation process elicited by the VSD domain is reversible by the changing membrane potential, long-lasting membrane depolarization also closes the channels, a process known as inactivation. In Kv channels, two well-defined inactivation processes have been identified, “N-type” inactivation and “C-type” inactivation. N-type inactivation underlies a rapid inactivation process in *Shaker* channels, which operates through a ball-and-chain mechanism in which one of the cytoplasmic amino terminal domains of the four protomers binds to the inner mouth of the pore (CC) and prevents ions from entering (2, 3), and is sensitive to intracellular blockers such as tetraethylammonium (TEA) and its derivatives (4). C-type inactivation is a slower process than N-type inactivation and is not inhibited by intracellular TEA (4) or truncation of the N terminal domain (2), unlike N-type inactivation, indicating that these two types of inactivation use different molecular mechanisms. C-type inactivation is primarily caused by conformational changes around the SF and/or outer mouth of the channels (4–6), and this conformational change occurs collectively among 4 protomers (7–9). The extracellular cations and TEA slow C-type inactivation (4, 10–12), suggesting a foot-in-the-door mechanism in which the binding of cations or TEA to extracellular binding sites prevents the conformational transition of the channel to the inactivated state. Extracellular K^+^ ions also modulate the recovery from C-type inactivation in Kv channels (13). Constriction of the SF underlying C-type inactivation occurs in a limited area near the external mouth (1, 14, 15). Cuello et al. suggested the molecular basis for C-type inactivation by solving the structures of the KcsA channel, which has no VSD, trapped in a series of partially open conformations of the SF (16). The backbone reorientation in the SF associated with inner gate opening leads to the loss of ion-binding sites in the SF and shortens the G77 diagonal Cα-Cα distances in the C-type inactivated state. Their group also detected interactions involving W67, which is equivalent to W434 in *Shaker* (italic font for *Shaker* numbering, e.g., *W434* will be subsequently used), G71, and D80 govern the rate and extent of C-type inactivation (17, 18). A point mutation in residue *W434* of the *Shaker* channel, corresponding to W366 of Kv1.2 or W67 of KcsA to a phenylalanine, results in C-type inactivation of the channels under a depolarizing membrane potential. They demonstrated the gating current resulting from structural rearrangement of the VSD (channel opening), without conduction of K^+^ ions (*W434F*) or rapidly entering a C-type inactivated state (W366F of Kv1.2) (19, 20). *W434* and *W435* compose the aromatic cuff and interact with *Y445* in the SF in the *Shaker* channel (21), affecting the conformational stability of the SF. Hoshi and Armstrong investigated the effect of the W434F mutation on the protein structure by homology modeling based on the crystal structure of Kv1.2-2.1 chimera and proposed that pore dilation rather than pore constriction causes C-type inactivation in Kv channels (22). Recently Conti et al. showed that VSD-to-pore coupling leads to an increased diameter of the outer mouth and loss of ion-binding sites (23). Narrowing of the intracellular part of the SF was also observed in their study.

To evaluate the detailed conformational changes in the SF in C-type inactivation and clarify their mechanism, we performed molecular dynamics (MD) simulations of the wild-type (WT) and W366F mutant of Kv1.2-2.1 chimera channel (pore domain) and compared their SF structures. The W366F mutant in our simulation showed nearly no conductance and remarkable structural differences in the carbonyl oxygen atoms of V375 and G376 residues compared to in the WT (G376 corresponds to G77 in KcsA.) Another difference was the coordination pattern of K^+^ ions in the SF. The two-ion state, in which S3 and S4 of the SF (shown in Fig. 1C) were occupied by K^+^ ions, was dominant in the W366F mutant, while the 3-ion state was dominant in the WT. The result of three-dimensional reference interaction site model (3D-RISM) analysis (24–26) for the WT and W366F mutant showed different probability distributions for K^+^ ions, suggesting that this 2-ion state in the W366F mutant (or difference in coordination number of K^+^ ions) is caused by structural differences of the SF in the W366F mutant. Furthermore, this conformational transition may occur depending on the voltage. We showed that the change in the relative orientations of the oxygen atoms in the SF leads to trapping of K^+^ ions in the SF and prevents its permeation.

**Figure. 1.**
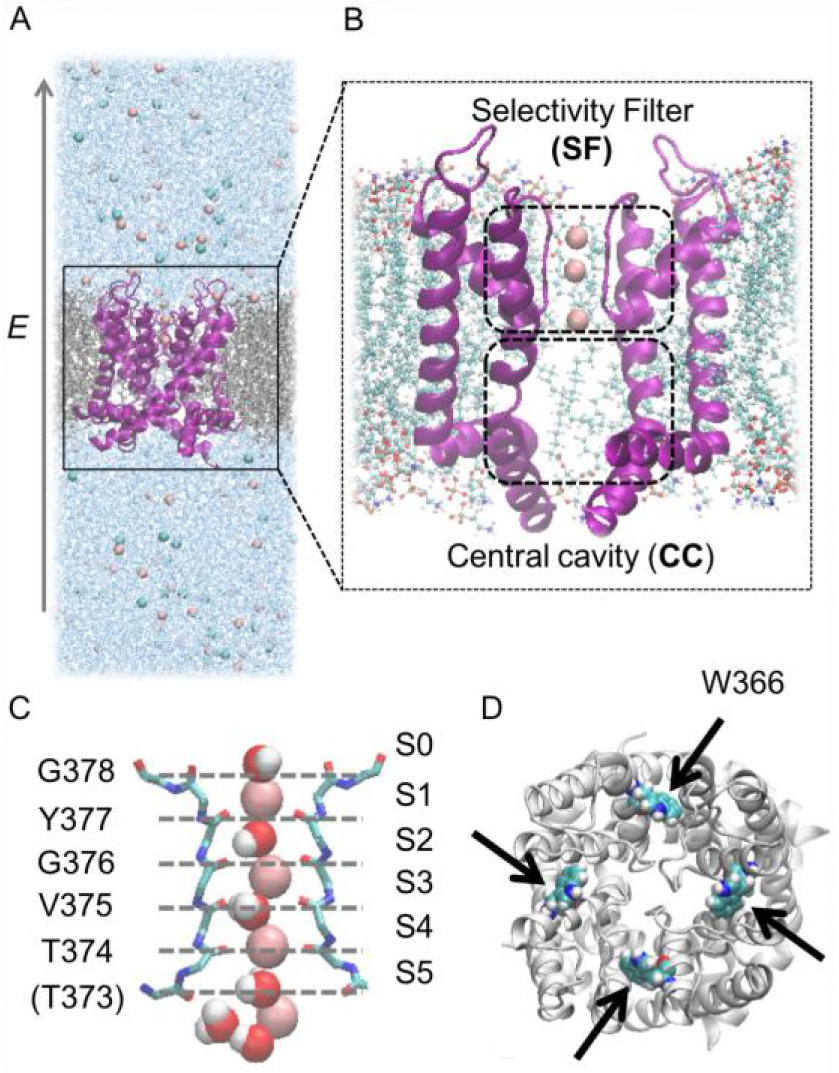
System of MD simulation. (A) Whole system of MD simulation. The system includes a pore domain of the potassium channel shown in purple NewCartoon, lipid bilayer (POPE molecules) in gray licorice, water molecules in blue points, and ions (K^+^ and Cl^−^) as pink and cyan particles. An electric field along the positive z-axis is applied to all atoms. (B) The enlarged view of a pore domain. The front and back protomers are not displayed for clarity. Ions enter the central cavity from the intracellular side and permeate through the SF in single file under the depolarization. (C) Structure of SF. The backbone heavy atoms of five residues (T374–G378) composing SF are shown in licorice. The front and back protomers were removed for clarity. Pink, red, and white particles represent K^+^, oxygen, and hydrogen atoms, respectively. (D) Pore domain viewed from the extracellular side. The mutated residues W366 are shown as space-filled spheres and indicated by black arrows.

## MATERIALS and METHODS

### System preparation

We used the X-ray structure of the Kv1.2-2.1 chimera channel (PDB ID: 2R9R) (27) as an initial structure for MD simulations. For W366F mutant simulation, the side-chains of the mutated residues were modeled by the AmberTools tLeaP program (28). Each system was solvated in a rectangular box with dimensions of approximately 70 × 70 × 180 Å^3^ (Fig. 1A); the length of the systems along the z-axis was increased enough to decrease the effect of the periodic boundary condition after a protein was embedded in a lipid bilayer consisting of 122 molecules of 1-palmitoyl-2-oleoyl-phosphatidylethanolamine (POPE). A lipid bilayer was constructed as described previously by Kasahara et al. (29). K^+^ and Cl^−^ ions were added to reach an approximate salt concentration of 340 mM. In this process, 3 K^+^ ions were retained in the SF, as in the X-ray structure (Fig. 1B). The SF consists of T374, V375, G376, Y377, and G378 residues of the four protomers, and the K^+^ coordination sites S0 to S5 were approximately defined by the carbonyl oxygen atoms of T373 and these five residues (Fig. 1C).

### Simulation details

All simulations were performed with GROMACS software version 4.5.3 (30). Simulations were carried out with the periodic boundary condition, and electrostatic interactions were treated with the particle mesh Ewald method (31, 32). CHARMM27 force field (CHARMM22/CMAP) (33–36) was applied for amino acids and lipid molecules, Jong and Cheatham ion parameters (37) for ions, and the TIP3P model (38) for water molecules. All bonds were constrained by the LINCS algorithm (39). We performed energy minimization of the whole system, and then gradually increased the system’s temperature using the Nose-Hoover thermostat (40, 41) with a position restraint for heavy atoms in the protein. After 10-ns equilibration, 300-ns production runs were performed with NPT ensemble, in which the system was controlled to 310 K and 1 bar by a Nose-Hoover thermostat and Parrinello-Rahman barostat (42, 43), respectively. For the mutant protein, a position restraint was used for the first half of the equilibration. The electrostatic field was applied along the z-axis for the whole system in the simulations with membrane potential. For an electric field of 0.004 V/Å, the voltage applied for the whole system was calculated to be approximately 700 mV based on the following formula: *V* = *EL* (*V*: voltage over the entire system, *E*: electric field, and *L*: length of a system along the z-axis), and this value is equivalent to the difference in electrostatic potential across the membrane as described previously (44, 45).

### Analysis by 3D-RISM

Ten snapshots with values of PC1 (first eigenvector) obtained from the principal component analysis (PCA) of values >0.25 or <-0.25 were sampled from the trajectories of the WT and W366F mutant, respectively. The lipid and solvent molecules were removed from the system, and a channel protein was considered as a solute in the electrolyte solution (KCl) at infinite dilution. The concentrations of K^+^ and Cl^−^ were set to 350 mM and the density of water molecules was set to 53.8 M. The Kovalenko-Hirata (KH) closure (24, 46) was used. The probability distributions were averaged over 10 snapshots. The number of grid points in the 3D-RISM-KH calculations was chosen as 256^3^ with 0.5-Å spacing. All 3D-RISM-KH calculations were conducted using in-house 3D-RISM codes (47, 48).

## RESULTS and DISCUSSION

### Stability of the systems

The time courses of root mean square deviation (RMSD) values were plotted in Fig. 2. We calculated RMSD values for two groups, Cα atoms in the whole protein (Fig. 2A) and carbonyl oxygen atoms in the SF (Fig. 2B), by fitting the atoms in each group to the initial structure by rotation and translation (least squares fitting). As shown in Fig. 2A, the mean values of RMSD for the whole protein were 1.5 Å for the WT and 1.6 Å for the W366F mutant, suggesting that the whole structures of the proteins were stable during both simulations. However, the RMSD values of local conformations of the SF in the mutant system (red line in Fig. 2B) were slightly higher than those in the WT system (black line in Fig. 2B), and locally showed lower values (close to values of WT) of approximately 60 ns. As described below, K^+^ ions seldom permeated through the W366F mutant, and conformations close to and distant from the initial structure represented the active (permeating) and inactive (non-permeating) states, respectively, of the SF in the mutant channel. Such fluctuations were not observed in the RMSD values of oxygen atoms of SF with least squares fitting of the Cα atoms of the whole protein (data not shown), suggesting a local conformational transition in the SF. Therefore, we describe the correlation between the local protein conformation and K^+^ ions in the SF (distributions and dynamics) in the following subsections.

**Figure. 2.**
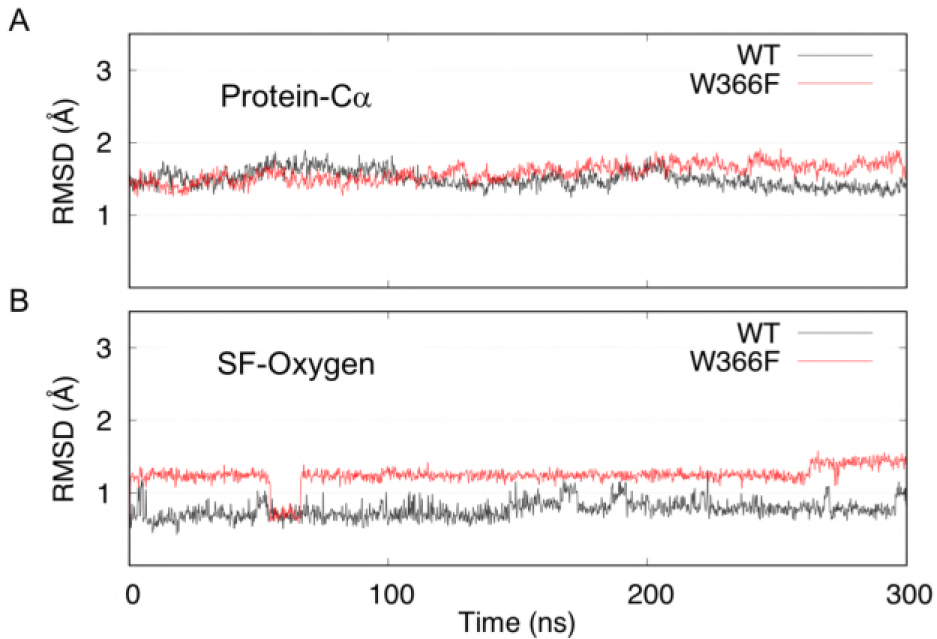
Time sequences of RMSD values. Time sequences of RMSD values of Cα atoms in a whole protein (A) and carbonyl oxygen atoms in SF (B). RMSD values were calculated after fitting the atoms in each group to the reference structures (initial structures) and plotted every 200 ps. The black and red lines represent WT and W366F mutant, respectively.

### Distributions of K^+^ ions and water molecules

To analyze the effect of the local conformational change in the SF in the W366F mutant on ion permeation, we compared the distributions of K^+^ ions and water molecules along the z-axis between the WT and W366F mutant. The numbers of K^+^ ions and water molecules in a cylinder with a radius of 5 Å centered on the channel pore (Fig. 3A) were counted for each x-y slice with a 1-Å thickness (Fig. 3B). Regions of the z coordinate of ~87 to ~102 and ~68 to ~87 were equivalent to the SF and CC, respectively. While the numbers of K^+^ ions and water molecules in bulk regions were nearly constant in both systems, some peaks were observed in the SF and CC. The location and magnitude of these peaks differed between the WT and W366F mutant, and we compared the distributions of each molecule (K^+^ ion or water molecule) in the SF region between them (Fig. 3C). (The peak around the entrance of the SF observed in the W366F mutant will be discussed later.)

**Figure. 3.**
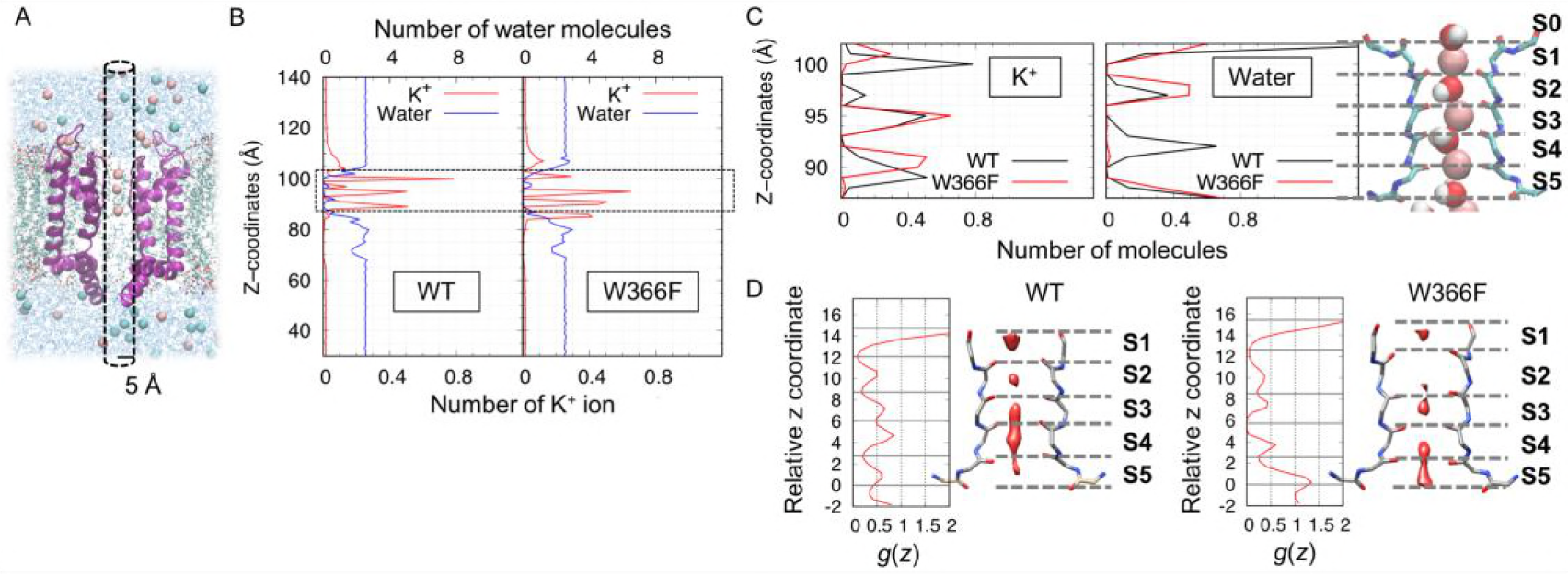
Distribution of K^+^ ions and water molecules. (A) Schematic diagram of the region in which the number of K^+^ ions or water molecules was counted. The x- and y-coordinates averaged over the 20-carbonyl oxygen atoms in SF were used as the center of a cylinder for each system. (B) Distribution of K^+^ ions and water molecules. The horizontal axis represents the number of K^+^ ions (lower scale) or water molecules (upper scale) in a cylinder with the radius of 5 Å averaged over each 300-ns simulation, and the vertical axis represents the z-coordinates. (C) Comparisons of distributions of K^+^ ions (left panel) or water molecules (right panel) in the SF region (shown in the dashed square in B) between WT and W366F mutant. The right figure represents the schematic diagram of SF. The expression is same as that in Fig. 1B. For K^+^ ions, 3 large peaks were observed around S1, S3, and S5 in WT, while there were 2 large peaks around S3 and S4 and a small peak around S1 in the W366F mutant. For water molecules, a peak around S4 disappeared in the W366F mutant. (D) Distributions of K^+^ ions averaged over 10 snapshots sampled from the WT (left panel) and W366F mutant (right panel) trajectories. In the right figure in each panel, the regions with high probability distributions of K^+^ ions (*g*(*r*) > 15.5) were shown as a red surface on each representative structure. The main-chain atoms of SF residues (T373–G378) are represented as sticks (only chains 1 and 2 are shown). The left graph in each panel shows the average probabilities over the xy plane (disc with a radius of 5 Å) along the z axis; 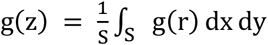, where *S* is the area of a disc. The gray lines represent the relative z coordinates of the oxygen atoms of SF residues (T373–G378) averaged over 10 snapshots: regions of S1–S5. The vertical axis represents relative values based on the z coordinate of the carbonyl oxygen atoms of T373: distance along the z axis from T373.

With respect to the distributions of K^+^ ions (left panel of Fig. 3C), there were three major peaks (around the S1, S3, and S5 regions) and a minor peak (S4) in the WT, whereas two major peaks (S3 and S4 regions) and a minor peak (outer mouth of SF) were observed in the W366F mutant. The major peaks were distributed at regular intervals in the WT (S1, S3, and S5). In contrast, in the W366F mutant, two major peaks were in the lower half of the SF in close proximity with each other (S3 and S4), and the minor peak was in the outer mouth of the SF, slightly distant from the major peaks, indicating the dissociation of K^+^ ions from S1 in the mutant system. This result corresponds to those of a previous study by Conti et al. (23). With respect to water molecules (right panel of Fig. 3C), the peak around S4 was not observed in the W366F mutant. This may be because the two K^+^ ions were located close to each other (S3 and S4) and a water molecule could not fit in the space between them.

For the molecular mechanism of C-type inactivation, we hypothesized that the local conformational change in SF led to a change in pore occupancy by K^+^ ions as observed in Figure 3C and the restriction of ion permeation. To evaluate this hypothesis, we analyzed the probability distributions of K^+^ ions in SF of the WT and W366F mutant by using the 3D-RISM theory to examine whether these two proteins have separate local conformations favoring different ion occupancy patterns with each other. Ten structures were sampled in each system as described in the Materials and Methods section, probability distributions were calculated for each structure, and then they were averaged over 10 structures in each system. The right figure in each panel of Fig. 3D shows the surface of probability distribution of K^+^ ions, *g*(*r*) of larger than 15.5: the region with high probability. In the WT, four large peaks and a small peak were observed in regions S1 through S4 and S5, respectively, and located in nearly the same interval, indicating that K^+^ ions can occupy all coordination sites in the SF. In contrast, in the W366F mutant, three large peaks and a small peak in regions S3 through S5 and S1, respectively, were observed and there was no peak in S2, suggesting that K^+^ ions are trapped around the entrance of the SF and that K^+^ ions around S1 are dissociated. The average probabilities over the xy plane (disc with a radius of 5 Å) along the z axis were also calculated (left graph in each panel of Fig. 3D). The gray lines in the graph are the average z coordinates of each residue and separate the SF into S1–S5. In the W366F mutant, the average probability distribution at S2 became 0.74-fold smaller than that of the WT. In addition, the probability distributions at the region between S1 and S2 and those between S3 and S4 were nearly 0. Considering the three-dimensional probability distribution of K^+^, these results suggest that the transition between S4 and S3 is restricted in the inactivated state. Figure 3D also shows that the vertical size of the SF changed in the W366F mutant (will be discussed in detail in a later subsection). The results of 3D-RISM analysis confirmed that the local conformational difference alters the pore occupancy by K^+^ ions and restricts ion permeation. We next analyzed the detailed difference in the SF conformation between the WT and W366F mutant.

### Structural differences between WT and W366F mutant protein

To detect conformational differences between the SF of the WT and W366F mutant we performed PCA for the coordinates of mainchain atoms in the SF (4 atoms × 5 residues × 4 protomers = 80 atoms). The rotational and translational motion was eliminated by fitting all structures to the average structure by referring to the backbone carbonyl oxygen atoms of residues in the SF (1 atom × 5 residues × 4 protomers = 20 atoms). A two-dimensional projection of the trajectories onto the PC1-PC2 space is shown in Fig. 4A. The contribution ratios of PC1 and PC2 were 41.7% and 6.6%, respectively. The SF mainchain conformations were roughly divided into 2 clusters by the PC1 (first eigenvector). Nearly all conformations from WT simulations showed positive PC1 values. In contrast, in the W366F mutant, most values clustered on negative PC1 values and some overlapped with those of WT, suggesting conformational transitions between the ‘open’ and ‘closed’ state occurring in the trajectory of the W366F mutant. Figure 4B shows the time course of PC1 values of both systems. The transition of PC1 values occurred at approximately 60 ns in the W366F mutant, corresponding to the local conformational transition suggested from the RMSD values (Fig. 2B).

**Figure. 4.**
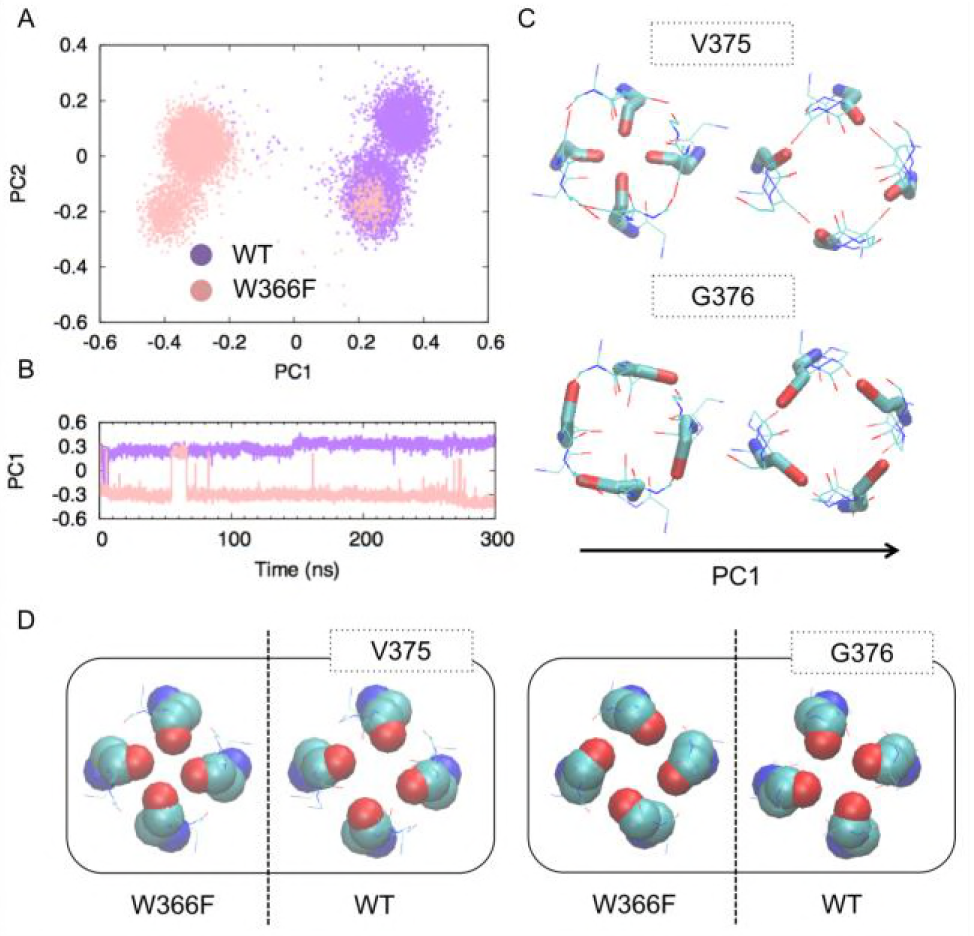
Results of PCA. (A) The projection of trajectories onto the first and second principal component axis (PC1 and PC2 values). PCA was applied for the Cartesian coordinates of the mainchain atoms of SF in the combined trajectory of WT and W366F mutant. (B) Time sequence of PC1 values of WT (purple line) and W366F mutant (pink line). PC1 values of W366F mutant were under 0 for nearly the whole simulation. The region where the PC1 values of W366F mutant overlap with those of WT (around 60 ns) corresponds to the area where the conformational transitions in SF occurred (Fig. 2B). (C) Schematic diagram of motion along the first eigen vector of V375 (upper) and G376 (lower) viewed from the extracellular side. The average structure of two trajectories was moved toward the negative or positive direction along the first eigenvector: PC1 = −1.0 (left) or +1.0 (right). (D) Typical structures (structures with the PC1 value averaged over each trajectory) viewed from the extracellular side. (PC1 = 0.292 for WT, −0.292 for W366F mutant). Mainchain heavy atoms of V375 (left panel) and G376 (right panel) residues of WT (right) and W366F mutant (left) are shown as a space-filling model.

The characteristic motions in the first eigenvector are represented in Fig. 4C. An increase in PC1 represents mainly a motion in which the carbonyl oxygen atoms of V375 move away from each other (Fig. 4C top) as well as a twist motion of the carbonyl oxygen atoms of G376 (Fig. 4C bottom). Thus, the decrease in PC1 from the WT to the W366F mutant leads to constriction of the SF or narrowing of the pore. Typical conformations of V375 and G376 of the WT and W366F mutant system, for which the PC1 values were averaged over each trajectory, are shown in Fig. 4D. Although we also performed PCA using not only the mainchain atoms of the SF, but also the whole protein structure (all Cα atoms), no conformational property characterizing W366F mutant was detected, suggesting that conformational changes in the SF are important for restricting ion permeation. In the next subsection, we will focus on the correlation between ion permeations and these local conformational changes.

### Correlation between ion permeation and SF conformation

The trajectories of K^+^ ions projected onto the z-axis in the WT and W366F mutant simulations are shown in the lower panels of Figs. 5A–B, respectively. Each line represents the time course of a position along the z-axis (z coordinates) of each K^+^ ion. In the WT, a 3- ion state, in which K^+^ ions occupied the S1, S3, and S5 sites, was observed most frequently, whereas in the W366F mutant, a 2-ion state, in which K^+^ ions occupied S3 and S4, was observed in nearly the entire simulation. K^+^ ions also remained around the entrance of SF over a long period in the W366F mutant. These results agree with the distributions of K^+^ ions described in the above subsection; a water molecule intervened mostly between two neighboring K^+^ ions in the WT but was not observed between the K^+^ ions occupying the S3 and S4 sites in the W366F mutant (Fig. 3C). The ions around the entrance of SF showed large fluctuations in CC (green and gray lines in Fig. 5B), suggesting that they were less stable than the two ions at S3 and S4.

**Figure. 5.**
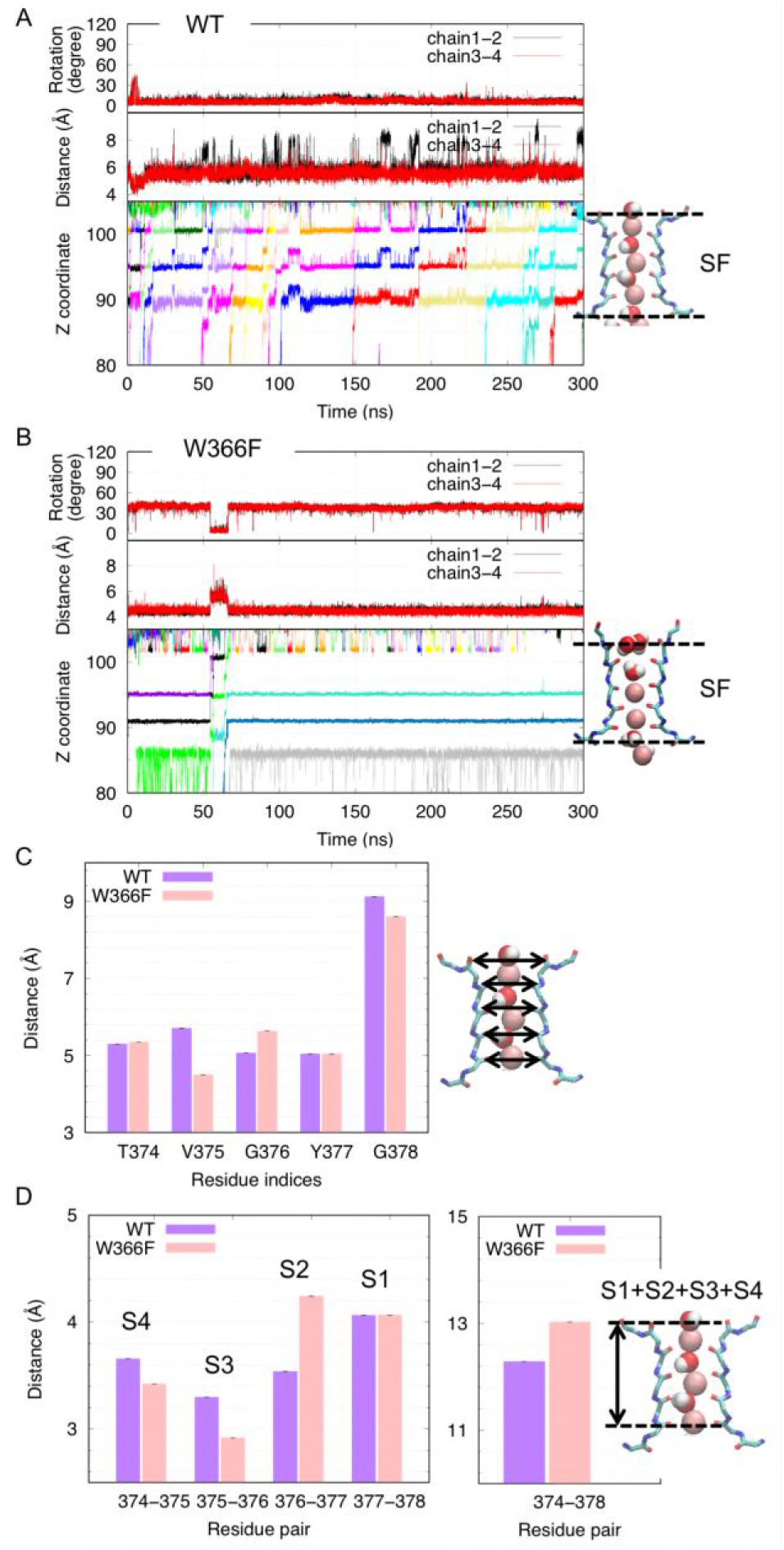
Correlation between ion permeation and SF conformation. (A–B) Time sequences of z-coordinates of each K^+^ ion (lower panel), distances between the backbone carbonyl oxygen atoms of V375 (middle panel), and rotation angle of the backbone carbonyl oxygen atoms of G376 (upper panel) of WT and W366F mutant, respectively. Z-coordinates (unit: Å) of ~90–105 and ~70–90 in the lower panel correspond the SF and central cavity regions, respectively. Distances between oxygen atoms of V375 in the middle panel were calculated for the pairs of chains 1 and 2 (black lines) and 3 and 4 (red lines). (C) Vertical size of the pore. The distances between the backbone carbonyl oxygen atoms of adjacent residues (vertical size of each region from S1–S4) and both terminal residues were averaged over time and for 4 protomers. The vertical bars on each box represent the standard error of the mean.

As shown in Fig. 5B, ion permeation, defined by release of a K^+^ ion from SF, was strongly reduced in the W366F mutant compared to in the WT. K^+^ permeation events were observed 19 and 3 times during the 300-ns simulations of the WT and W366F mutant, respectively. All ion permeation events in the W366F mutant occurred at approximately 60 ns, when local conformational transitions from the ‘closed’ to ‘open’ state occurred in the SF (Fig. 4). This reveals a correlation between ion permeation and the local conformation of the SF. In the following subsections, we describe the detailed conformational difference in the SF between the ‘active’ and ‘inactive’ states.

### Distances between backbone carbonyl oxygen atoms composing SF

The conformational change in V375 was one of the most remarkable motions corresponding to the first eigenvector of PCA (Fig. 4C). To visualize the conformational characteristics of SF of the W366F mutant, pore sizes at 5 sites (residues composing SF; T374, V375, G376, Y377, and G378) were analyzed. We calculated the distances between the backbone carbonyl oxygen atoms of each residue in protomers facing each other: chains 1 and 2 and chains 3 and 4. As expected from the PCA (Fig. 4) we found a correlation between the pore size at V375 residues in the middle of S2 and S3 (middle panels of Figs. 5A–B) and ion permeation (lower panels of Figs. 5A–B). In the WT, the distances between V375 of both pairs fluctuated around 5.5 Å. In contrast, the distances were stable around 4.5 Å in the W366F mutant, except for a temporary and concordant increase in the two distances to ~5.5 Å at the 50–70-ns region of the trajectory, in which momentary ion permeations were observed. Although in residues other than V375, no remarkable transitions of the atom-atom distance correlating with ion permeation events were detected (data not shown), the pore sizes at G376 and G378 (average distances between oxygen atoms) averaged over the trajectories were slightly different between the 2 systems (Fig. S1 in the Supporting Material). The conformational difference in G376 is described in the next subsection.

The vertical length of the pore also differed between the WT and W366F mutant. We calculated the distances between the carbonyl oxygen atoms of adjacent residues: T374 and V375, V375 and G376, G376 and Y377, and Y377 and G378, and the distance between terminal residues T374 and G378 for each chain. These results were averaged over the protomers. As shown in Fig. 5C, the whole length of the pore (distance between T374 and G378) of the mutant proteins was larger than that of the WT. As suggested from the conformational changes of V375 and G376, the sizes of the three coordination sites flanking V375 and/or G376 were significantly changed; the S2 (G376-Y377) region was especially elongated and the S3 (G374-V375) and S4 (V375-G376) regions were shortened.

### Rotation of backbone carbonyl oxygen atoms of G376

To quantify the twist motion of the carbonyl group of G376, we calculated the angle between the vector connecting the backbone carbonyl oxygen atoms of G376 of the two protomers facing each other (from chains 1 to 2) in each snapshot and the same vector in the reference structure. The same calculation was carried out for another pair of protomers (from chains 3 to 4). The initial structure of the WT simulation was used as the reference structure to evaluate the rotation of both the WT and W366F mutant, and therefore the angle of 0 represents the same conformation as the WT initial structure. As shown in the upper panels in Figs. 5A–B, rotation angles fluctuated around ~0° in the WT and ~40° in the W366F mutant. A temporary transition from ~40° to ~0° was observed in the mutant protein at 50–70 ns of the simulation time, which correlated with the recovery of the channel and pore size at V375 (Fig. 5B), suggesting that this conformational transition is related to ion permeability.

### Voltage dependence of the SF structural change

In the trajectory of the W366F mutant, the transition from the active state to the inactivate state occurred immediately after equilibration. To analyze the voltage dependence of the structural change of the SF, we performed a 300-ns simulation of the W366 mutant with no applied voltage. The initial structure was the same as that in the simulation with an applied voltage. The time course of the z coordinates of K^+^ ions and projection onto PC1 obtained from the aforementioned PCA of the WT and W366F trajectories are shown in the lower and upper panels of Fig. 6A, respectively. The distribution of PC1 values was nearly the same as that obtained from WT simulation (Fig. 4B) (although PC1 values sometimes became negative, a state with a negative PC1 value was not long-lasting), revealing that inactivation of the channel was almost not observed in this simulation. The three-ion state (S1, S3, and S5 were occupied) was maintained during the nearly whole simulation, but an ion closest to the CC (black line in Fig. 6A, around S5) was not stable and fluctuated between S5 and S4.

**Figure. 6.**
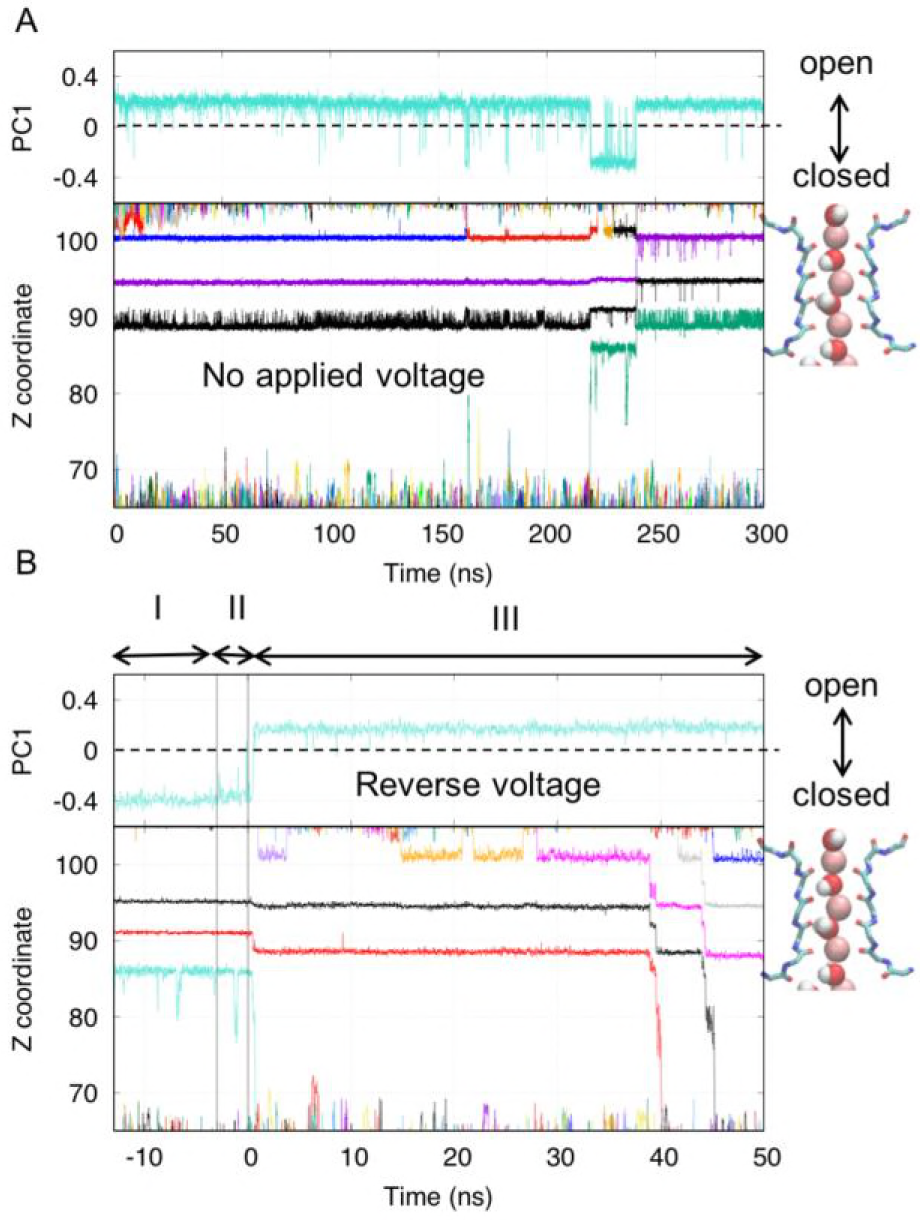
Voltage dependence of the SF structural change. (A) Result of a 300-ns simulation with no applied voltage. The lower and upper panel show time sequences of z-coordinates of each K^+^ ion and projection onto the first eigenvector obtained from PCA of SF conformations of WT and W366F mutant (PC1 values), respectively. An ion permeation event associated with a conformational transition between ‘open’ and ‘closed’ was observed. There was no large conformational change in SF (the distribution of PC1 values were same as that of WT except for the region ~220–240 ns. (B) Result of a 50-ns simulation with reverse voltage. The region I represents the last 10 ns (from 290–300 ns) of the trajectory of W366F mutant with an applied voltage of 0.004 V/Å. The applied voltage gradually decreased from 0.004 to −0.004 V/Å during region II, and then was fixed at −0.004 V/Å in region III. The ion state and PC1 values drastically changed at approximately 0.5–0.6 ns.

Additionally, we performed a 50-ns simulation of an inactivated channel with the reverse voltage (E = −0.004 V/Å). The final structure (300 ns) of the trajectory of the W366F mutant simulation with the voltage of 0.004 V/Å (inactivated state) was used as the initial structure, and the applied voltage gradually changed from 0.004 V/Å to −0.004 V/Å during 3- ns simulation (region II: from −3 to 0 ns in Fig. 6B). The upper panel of Fig. 6B showed that the conformational transition in the SF occurred immediately after the change in voltage (around 0.5–0.6 ns). The ion state also changed at the same time (from S3-S4 to S3-S5). Two ions passed through the pore from the outside to the inside. Recovery was not observed in the 100-ns simulation with the same initial structure and no applied voltage (0.0 V/Å) (data not shown), suggesting that some energy (or time) is required for recovery from inactivation.

The results of these two simulations indicate the voltage dependence of the conformational transition of SF between the inactivated state and activated state. These simulations explain the experimental result of fast inactivation of the W366F mutant upon membrane depolarization.

### Conformational difference around the outer mouth of SF between the WT and W366F mutant

Previous studies revealed that the exposure of residues in the extracellular region increase upon C-type inactivation (5, 49), suggesting that conformational changes associated with C-type inactivation also occur in the extracellular region of the pore. Solvent-accessible surface areas (SASA) of M380 (*M448*) residues (total value of 4 residues in each chain) were calculated for every 50 snapshots (interval of 1 ns). The averaged value of SASA over 300-ns trajectories of the W366F mutant (210.3 ± 25.7 Å^2^) was larger than that of the WT (153.4 ± 29.0 Å^2^). We also performed PCA for the coordinates of the heavy atoms of M380 to detect conformational differences causing the increase in SASA values. The scatterplot in Fig. 7A shows the 2-dimensional projection of trajectories on the first and second eigenvectors (PC1 and PC2 values), and the histogram above this plot shows the distribution of PC1 values for each trajectory, which suggested that the PC1 values for M380 distinguish the conformations in the WT and W366F mutant trajectories. The contribution ratios of PC1 and PC2 for M380 were 29.9% and 13.6%, respectively. Figure 7B shows the mode of PC1 for M380. The upper panel (structure moving to the positive direction, large PC1 values) and lower panel (structure moving to the negative direction, small PC1 values) suggest that the side chains of 4 methionine residues move alternately along the first eigenvector. Remarkable differences between the WT and W366F mutant were not detected in the distances between the Cα atoms or sulfur atoms of M380 of chains 1 and 2 or 3 and 4, respectively (data not shown).

**Figure. 7.**
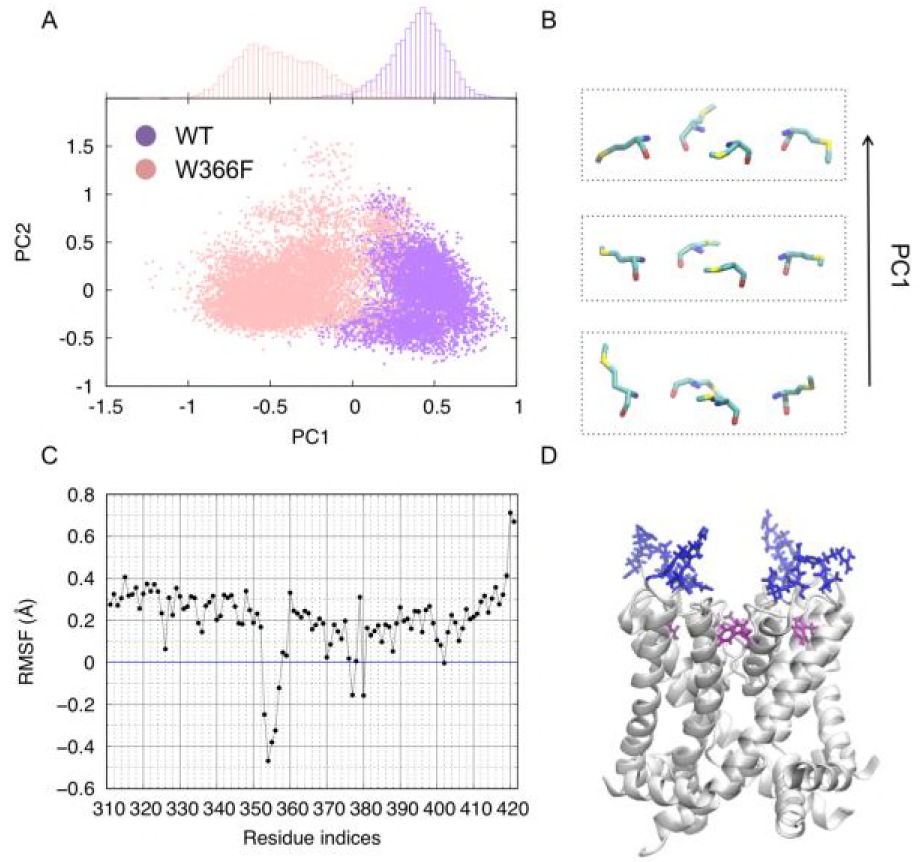
Conformational difference around the outer mouth of SF. (A) Two-dimensional projection of the results of PCA for M380 sidechain conformations. The purple and pink points represent WT and W366F mutant trajectories, respectively. Histograms on the graph show the distribution of PC1 values for each trajectory (purple: WT and pink: W366F). (B) Snapshots of structure morphing along the first eigenvector for M380. The middle panel represents the average structure and upper and lower panels show structures moved to the positive and negative directions from the average structure, respectively. (C) Differences in RMSF values between WT and mutants versus residue indices. The RMSF values of W366F were subtracted from those of WT. (D) Residues with negative ΔRMSF (displayed in blue licorice). Nearly all residues are in the extracellular region. The white NewCartoon represents the structure of the pore domain taken from the WT trajectory (snapshot at 100 ns simulation time), and the residues shown as purple licorice are W366.

To analyze the fluctuations of W366 or F366, we calculated the root mean square fluctuation (RMSF) values of protein heavy atoms by residues for 300-ns trajectories of each system. There was no large difference in the RMSF values of W366 or F366 among the systems: 1.28 and 1.44 Å for the WT and W366F mutant, respectively. The difference in RMSF values of each residue between the WT and W366F mutant is shown in Fig. 7C. The RMSF values in the WT were ~0.2–0.4 Å larger for most residues, but smaller for residues 353–357, 377, and 380 compared to those in the W366 mutant. Residues 353–357 and 380 were all in the extracellular region and 377 was in the SF (Fig. 7D), suggesting that fluctuations of residues around the extracellular mouth change upon conformational changes around the SF. Particularly, RMSF values of residue Y377 became large in one or two protomers in the W366F mutant.

## CONCLUSION

We performed MD simulations of the W366F mutant, which showed faster C-type inactivation, and observed the conformational difference in the SF from the ion-permeable and impermeable states. The results of PCA showed the characteristic conformational difference between SF structures of the WT and W366F mutant: distortion of the carbonyl oxygen atoms of G376 and pore narrowing at V375. This conformational change was voltage-dependent and prevented ion permeation, suggesting that this is the molecular basis of C-type inactivation. Considering the results of Conti et al. (23) in which the conformational changes enlarging the outer mouth of SF in addition to making the middle of SF narrow were observed, the structural change observed in this study may be part of a transition to C-type inactivation and the W366F mutation facilitate this conformational transition depending on the membrane potential. In addition to the structural difference, ion states differed between the WT and W366F mutant. While the 3-ion state in which S1, S3, and S5 of SF were occupied by K^+^ ions were important in the WT, the W366F mutant remained in the 2-ion state occupied by S3 and S4. In the W366F mutant, K^+^ ions in S1 dissociated, although the pore size at S1 did not greatly change. The ion distributions obtained from 3D-RISM analysis also showed that K^+^ ions are likely to be in the S3–S5 in the W366F mutant. Furthermore, additional 3D-RISM analysis for the W366F mutant in the active state (PC1 values > 0.25) indicated that proteins in the active state showed a similar distribution of K^+^ as the WT even in W366F mutant (Fig. S2), revealing that the ion state observed in the W366F mutant is caused by the conformational change of SF. In summary, the conformational change narrowing the middle of the SF occurred in the W366F mutant depending on the applied voltage, causing a loss of the ion-binding site around S1 and the stagnation of K^+^ ions in the S3 and S4 regions. These results observed in the W366F mutant simulation should be the molecular basis of C-type inactivation.

## AUTHOR CONTRIBUTIONS

HXK, KK, and MS conceptualized the study. HXK and NY carried out computational analyses and wrote the original draft of the manuscript. MS and KK contributed to reviewing and editing of the manuscript. KK supervised the study. All authors approved the final version of the manuscript and publication.

## ACKNOWLEDGEMENTS

This study was supported by MEXT Grant-in-Aid for Scientific Research on Innovative Areas “Integrative Multi-Level Systems Biology” and Platform Project for Supporting Drug Discovery and Life Science Research (Basis for Supporting Innovative Drug Discovery and Life Science Research (BINDS)) from AMED under Grant Number JP18am0101067. We thank RIKEN Advanced Center for Computing and Communication for providing computational resources of HOKUSAI. The authors declare that they have no conflicts of interest with the contents of this article.

